# MicroED Structures of HIV-1 Gag CTD-SP1 Reveal Binding Interactions with the Maturation Inhibitor Bevirimat

**DOI:** 10.1101/241182

**Authors:** Michael D. Purdy, Dan Shi, Jakub Chrustowicz, Johan Hattne, Tamir Gonen, Mark Yeager

**Author notes:** **Corresponding author:** Mark Yeager, M.D., Ph.D., Sheridan G. Snyder Translational Research Building, Rm 320, 480 Ray C. Hunt Drive, Charlottesville, VA 22908.

## Abstract

HIV-1 protease (PR) cleavage of the Gag polyprotein triggers the assembly of mature, infectious particles. Final cleavage of Gag occurs at the junction helix between the capsid protein CA and the SP1 spacer peptide. Here we used MicroED to delineate the binding interactions of the maturation inhibitor bevirimat (BVM) using very thin frozen-hydrated, three-dimensional microcrystals of a CTD-SP1 Gag construct with and without bound BVM. The 2.9-Å MicroED structure revealed that a single BVM molecule stabilizes the 6-helix bundle via both electrostatic interactions with the dimethysuccinyl moiety and hydrophobic interactions with the pentacyclic triterpenoid ring. These results provide insight into the mechanism of action of BVM and related maturation inhibitors that will inform further drug discovery efforts. This study also demonstrates the capabilities of MicroED for structure-based drug design.

**One-sentence summary:** The MicroED structure of HIV-1 Gag CTD-SP1 at 2.9 Å resolution reveals the molecular basis for binding of the maturation inhibitor bevirimat that will inform further drug discovery efforts.

---

HIV-1 Gag and Gag-Pol assemble and bind the plasma membrane of infected cells ^4^ The packing interactions between Gag hexamers induces positive membrane curvature, and engagement of cellular ESCRT complexes results in budding and release of immature HIV-1 virus particles comprised of the membrane-encased shell formed by the Gag lattice and the viral RNA. In the immature virion, Gag hexamers form a fissured, spherical shell with quasi-hexagonal symmetry on the inner leaflet of the viral envelope^5-7^. HIV-1 viral protease (PR) is liberated from Gag-Pol by autoproteolysis at a late stage of viral assembly, which ensures that Gag molecules are not processed before assembly of the lattice. All five Gag processing sites are essential for infectivity, and the sites are processed at different rates, in the order: SP1-NC > MA-CA ~ SP2-P6 > NC-SP2 > CA-SP1. The final, CA-SP1 cleavage triggers the dramatic morphological changes between immature and mature infectious HIV-1 particles that contain the characteristic conical capsid^8,9^. Inhibition of Gag cleavage effectively prevents HIV-1 replication, and two therapeutic strategies have been pursued: protease inhibitors that bind directly to PR, and maturation inhibitors, which interfere with PR cleavage of Gag. The first characterized maturation inhibitor was bevirimat (BVM), which acts by preventing PR cleavage between residues L363 and A364 in the junction helix formed by the C-terminal domain of CA and SP1 spacer peptide, thereby stabilizing the immature lattice^10 11^ However, the mechanism by which BVM stabilizes the immature lattice is unclear. Although BVM entered clinical trials, efficacy was thwarted by the presence of preexisting resistant viruses containing SP1 polymorphisms^12,13^. Second generation maturation inhibitors have improved resistance profiles^14,15^ but rational drug development has been impeded by a lack of high-resolution structural data on maturation inhibitor binding to Gag.

In 2016, compelling evidence for the predicted 6-helix bundle structure of the junction helices was provided by electron cryotomography and subtomogram averaging of ΔMA-Gag, non-enveloped virus-like particles (VLPs)^16^, X-ray crystallography of a Gag construct comprised of the C-terminal domain (CTD) of CA and SP1^17^, and solid-state △MR spectroscopy of △MA-Gag VLPs^18^. The cryoEM and X-ray structures revealed that the protease cleavage sites are sequestered on the interior of the 6-helix bundle. The cryoEM map also showed that a single molecule of the maturation inhibitor bevirimat (BVM) binds at the center of each 6-helix bundle. To achieve a resolution of 3.9 Å, 6-fold averaging was applied to the cryoEM map, which prevented characterization of the interactions between BVM and the CTD-SP1 junction helices.

Here we used MicroED^19-23^ to delineate the binding interactions of BVM using very thin frozen-hydrated, three-dimensional microcrystals of the CTD-SP1 Gag construct. The protein was expressed and purified as described previously^17^, and the crystallization conditions were adapted to grow microcrystals < 1 μm thick (as described in the Materials and Methods and **Supplementary Fig. 1**). We solved the structures of drug-free and BVM-bound CTD-SP1 by MicroED (i.e., electron diffraction from frozen-hydrated, three-dimensional microcrystals) to resolutions of 3.0 Å and 2.9 Å, respectively (Table S1), without applying 6-fold symmetry.

## RESULTS

The CTD-SP1 crystals were of the same form that was used to solve the drug-free X-ray crystallographic structure to 3.27 Å resolution^17^. The asymmetric unit contains one CTD-SP1 hexamer, and layers of the C2 crystals approximate the HIV-1 Gag immature lattice (**Fig. 1**). Molecular replacement phasing of the CTD-SP1-BVM MicroED data with the CTD X-ray structure (PDB 5I4T) resulted in a map that contains BVM density at the center of the 6-helix bundle formed by the CTD-SP1 junction helices (**Fig. 1d-f**). The density attributed to BVM is absent in our MicroED map of drug-free CTD-SP1 (**Fig. 1a-c**). The location of the single BVM molecule within the 6-helix bundle is consistent with a 3.9-Å map of HIV-1 Gag virus like particles (VLPs) determined by electron cryotomography and subtomogram averaging^16^.

**Figure 1.**
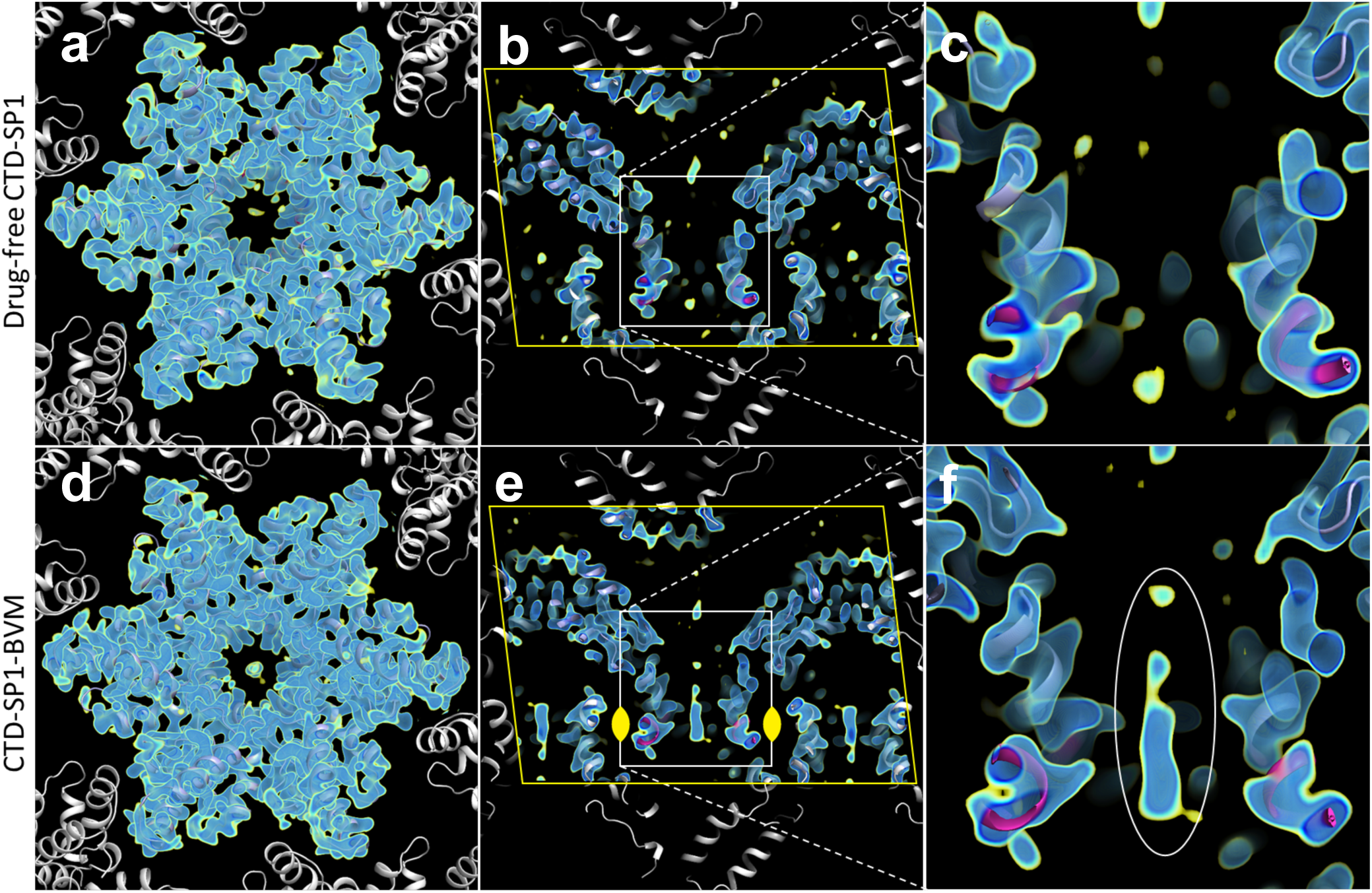
MicroED maps of drug-free (3.0 Å) and BVM-bound (2.9 Å) CTD-SP1. Maps include multiple contour levels from 1.2 σ (yellow) to 2.0 σ (dark blue). Indicated in **b** and **d** are crystallographic symmetry mates (white), unit cell edges (yellow line), and the 2-fold symmetry operator (yellow). **(a** and **d**) Views along the hexamer axis showing the CTD domain and packing that mimics the immature HIV-1 lattice. (**b** and **e**) Cutaway of the CTD-SP1 hexamers along the hexamer axis (unit cell boundary, yellow line). (**c** and **f**) Close-ups showing the center of the 6-helix bundles (CTD, slate blue and SP1, magenta), with BVM density present in **f** (white ellipse).

In the case of the cryoEM map of VLPs, the BVM density was a featureless ellipsoid due to 6-fold averaging required to achieve a resolution of 3.9 Å for the protein density. However, 6-fold averaging was not applied in the determination of the CTD-SP1-BVM MicroED map, and the BVM density is asymmetric with a dominant feature consistent with en face (**Fig. 2c**) and profile (**Fig. 2d**) views of the pentacyclic triterpenoid moiety of BVM. The presence of a unique mode of binding is presumably a consequence of the two-fold symmetric crystallographic environment of the binding site (**Fig. 1e**).

**Figure 2.**
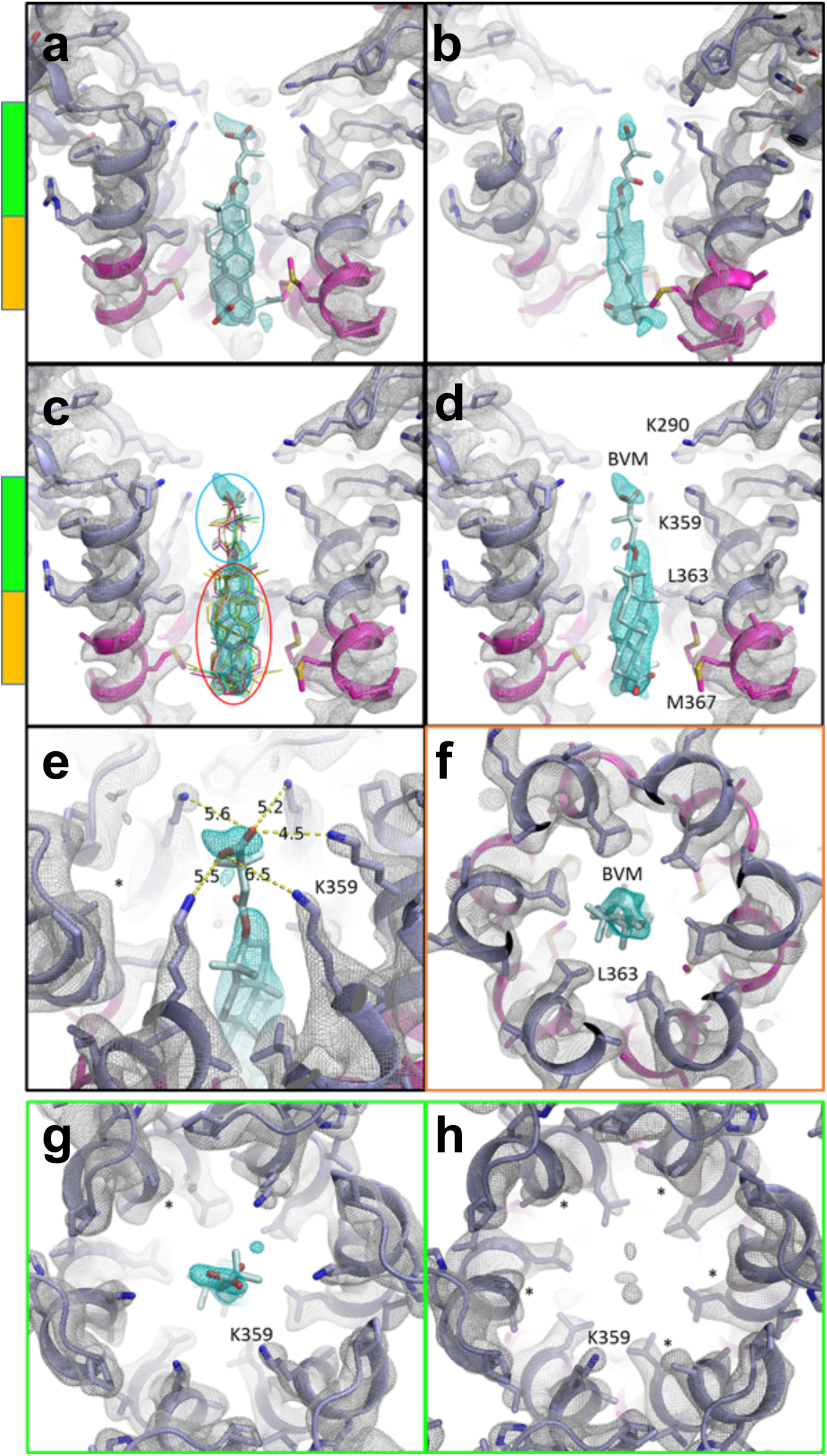
MicroED map of CTD-SP1-BVM suggests a mode of drug binding. CTD-SP1-BVM map contoured at 1.2 o above the mean (a single map is shown with protein density colored white and BVM density cyan). Green and orange bars indicate the axial positions shown in **f**, **g**, and **h**. *BVM was never included in the model used for map calculation*. (**a**) *En face* and (**b**), profile views of BVM density within the 6-helix bundle formed at the junction of CTD (slate blue) and SP1 (magenta). (**c**) Eleven BVM conformers^3^ docked into the map show that the rigid pentacyclic triterpenoid (red ellipse) fits well into the primary drug density, while the terminal carboxyl of each conformer fits within the satellite peak. Flexibility of the dimethylsuccinyl group (blue ellipse) is likely responsible for the break between the primary and satellite drug densities. (**d**) A single BVM conformer shows that drug binding spans the length of the junction helices and positions the dimethylsuccinyl carboxyl at the center of the K359 ring of lysines (**e**) The K359 side chain densities and B-factors vary amongst the protomers, supporting a preferred BVM binding pose with associated, specific K359 interactions. The side chain of one K359 (*) was not modeled due to a complete lack of density. (**f**) L363 of the CTD-SP1 protease cleavage site contributes the preponderance of hydrophobic interactions with BVM. The opposite axial orientation (“BVM-down”) leaves the satellite peak unoccupied and positions the C28 carboxyl of the pentacyclic triterpenoid distant from K359 **(Supplementary Fig. 2). (g)** Axial view of the BVM binding site showing density for 5 of 6 K359 side chains near the terminal BVM carboxylic acid, **(h)** Axial view of drug-free CTD-SP1 showing the absence of density for 5 of 6 K359 side chains.

The location of the pentacyclic triterpenoid adjacent to the junction helices clearly indicates that hydrophobic interactions are important for the mechanism of binding of BVM (**Fig. 3e,f**). However, the primary drug density of the ring structure does not define the axial orientation of the bound drug, i.e., whether the dimethylsuccinyl group is oriented toward (BVM-up) or away from (BVM-down) the CA domain (**Supplementary Fig. 2**). Regardless of the axial orientation, the hydrophobic interactions between BVM and Gag are confined to the junction helices. Specifically, L363 and M367 from each of the six protomers contribute the majority of the BVM binding interactions (**Fig. 2c, d and f**).

**Figure 3.**
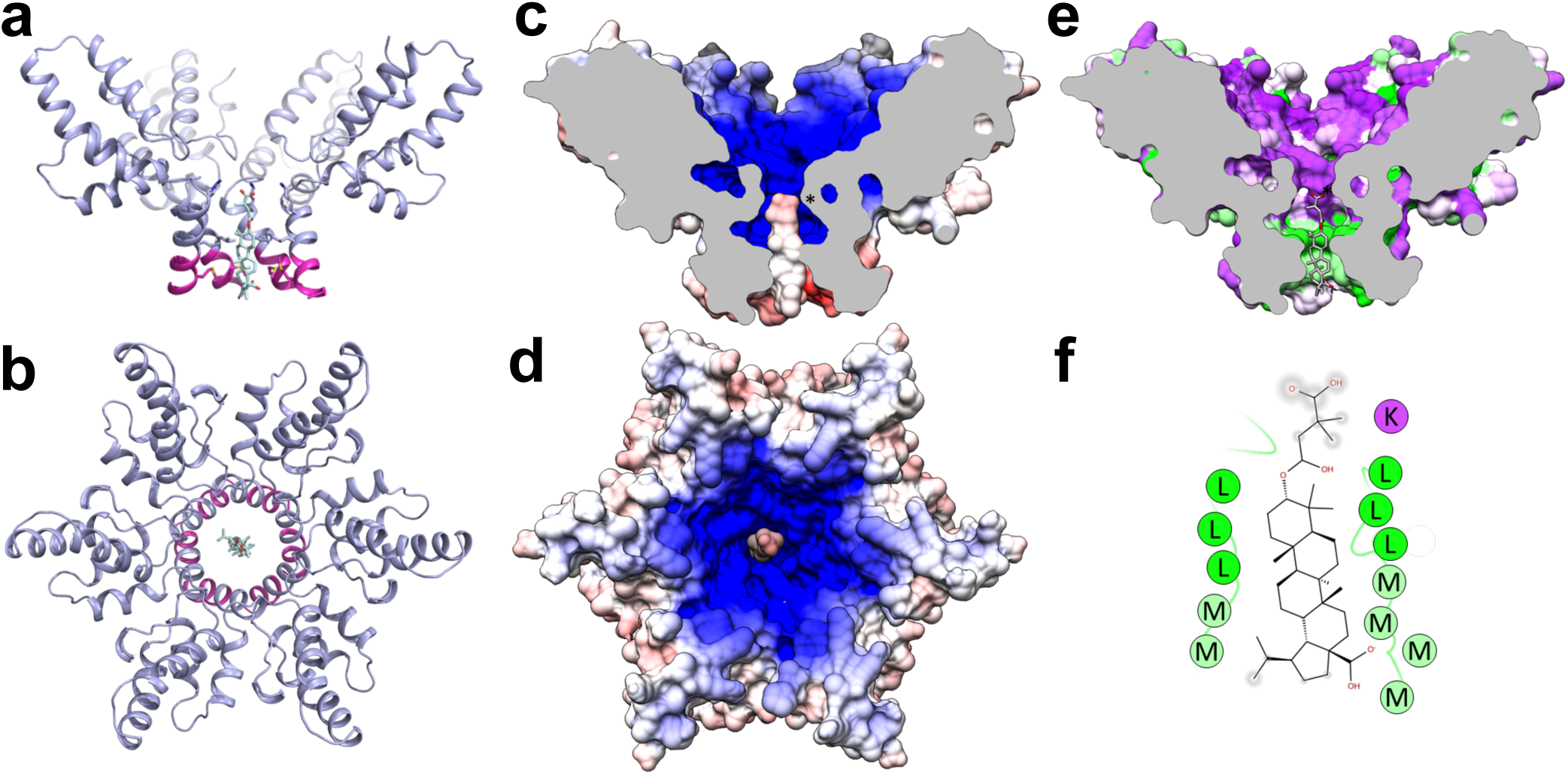
BVM binding to Gag is mediated by electrostatic and hydrophobic interactions. Ribbon model of the CTD (slate blue) and SP1 (magenta) 6-helix bundle with bound BVM, viewed from the side **a**, and top **b. (c** and **d**) CTD-SP1 and BVM electrostatic surface potentials show that the negative charge of the dimethylsuccinyl carboxyl resides at the center of the positive charge generated by the dual-ring of lysines (K290 and K359). **(e)** BVM binding also includes hydrophobic interactions between the junction helices of each Gag protomer and the pentacyclic triterpenoid of BVM (green, hydrophobic to purple, polar). **(f)** Schematic two-dimensional representation of BVM binding interactions showing hydrophobic (green) and polar (purple) residues. PR cleavage occurs at the L363 residues, which contribute the majority of the hydrophobic interactions with BVM.

The axial orientation of the BVM is suggested by a satellite density directed toward the CA domain, which we ascribe to the dimethylsuccinyl moiety (**Fig. 2a-e**). This BVM-up orientation places the carboxylic acid of the dimethylsuccinyl moiety in proximity with a ring of lysine residues (K359). Notably, the extent of the density of the K359 side chains varies between protomers in the BVM-bound map. In particular, the side chain density is better defined for the K359 residue that is closest to the density peak we ascribe to the dimethylsuccinyl (**Fig. 2e,g**). In contrast, density for the K359 side chain is absent in 5 of 6 protomers in the drug-free CTD-SP1 map (**Fig. 2h**), likely due to conformational flexibility. The presence of density for the dimethylsuccinyl carboxylate of BVM may seem surprising since acidic side chains frequently have diminished density in cryoEM maps^24^. However, proximity to the K359 ring of lysine likely neutralizes the Coloumb potential of the carboxylate, rendering it less susceptible to the nuances of electron scattering (for a detailed discussion see Supplementary Methods and **Supplementary Fig. 3**).

Refinement of the structure with BVM in the up orientation also supports our interpretation (**Supplementary Fig. 2a**), whereas refinement with BVM in the down orientation positions the C-28 carboxyl more than 7 Å away from the nearest K359 Nζ, and the dimethylsuccinyl extends out of the junction helix bundle (**Supplementary Fig. 2b,c**).

## DISCUSSION

BVM is known to act by stabilizing the immature Gag lattice^11^, and the BVM dimethsuccinyl ester is thought to contribute to the activity^15^. It is possible that BVM stabilizes Gag hexamers by electrostatically anchoring to K359 of the CA_CTD_ domain via the dimethylsuccinyl carboxylate (**Fig. 3c,d**) and binding together the junction helices via hydrophobic interactions between L363 and M367 from each protomer and the BVM pentacyclic triterpenoid moiety (**Fig. 3e,f**). This BVM binding mechanism could accommodate lattice and hexamer distortions due to the non-specific nature of the hydrophobic interactions and the electrostatic attraction with the six K359 residues, which is amplified by a second, coaxial ring of lysine residues (K290) (**Fig. 2c,d**).

This mechanism of BVM binding and Gag stabilization is also consistent with the structures of second-generation HIV-1 maturation inhibitors in which the dimethylsuccinyl ester is replaced with benzoic acid (**Supplementary Fig. 4**). C-3 benzoic acid substitution places a carboxyl in proximity to K359 while preserving the hydrophobic interactions between the junction helices and the pentacyclic triterpenoid. The fact that the C-3 benzoic acid substituted compound exhibited antiviral activity comparable to BVM and had no effect on the potency against the BVM Gag escape mutant V370A provides additional support for BVM binding with the succinic acid positioned in the vicinity of K359 and away from V370. In addition, the BVM-up orientation positions C-28 of BVM near M367, and C-28 derivatives that extend BVM analogs toward the C-terminal end of SP-1 (**Supplementary Fig. 5**) have the highest potency against HIV-1 with the Gag V370A polymorphism^13-15^. BVM analogs with C-28 extensions may provide additional, compensatory interactions at the C-terminal end of the 6-helix bundle that are sufficient to prevent protease access to the junction helix cleavage site.

We realize that our interpretation of the binding mechanism of BVM to the 6-helix bundle is based on comparison of MicroED structures with and without the drug. For instance, photoaffinity labeling of detergent-stripped VLPs suggested that BVM may bind in the down orientation ^25^. In the context of assembly of the immature hexagonal Gag lattice at the plasma membrane, it may be possible that other factors such as binding of viral RNA may influence the up-down distribution of BVM orientations. Likewise, the orientation may be influenced by the chemical properties of R-group substitutions.

Binding of a single BVM molecule to the Gag hexamer presents challenges for structure-based drug design, but this mode of binding is likely responsible for the efficacy of BVM and related maturation inhibitors that bind to Gag, despite structural heterogeneity inherent in the quasi-hexagonal immature lattice. The CTD-SP1-BVM structure provides additional insights into BVM binding and activity that will inform further development of HIV-1 maturation inhibitors.

The CTD-SP1-BVM structure also demonstrates the capability of MicroED for structure-based drug design and discovery even in challenging cases (i.e., low crystallographic symmetry and diffraction to moderate resolution). The increasing availability of electron microscopes and the ability to obtain microcrystals, either by screening under low dose in an electron microscope or generated by mechanical disruption of larger crystals^26^, make MicroED an attractive option for high-resolution macromolecular structure determination, even for proteins that are far too small for analysis by single-particle cryoEM methods.

## METHODS

The protein was expressed and purified as described previously^17^. The crystallization conditions were adapted to grow microcrystals, the data were collected and the structures were solved as described in the Supplementary Methods. Any additional supplementary display items, source data and associated references, are available in the online version of the paper.

## ACKNOWLEDGMENTS

This study was funded by U.S. National Institutes of Health grants to M.Y. (R01 GM066087 and P50 GM082545). We gratefully acknowledge Eric Freed for a critical reading of the manuscript. Some molecular graphics and analyses were performed with the UCSF Chimera package. Chimera is developed by the Resource for Biocomputing, Visualization, and Informatics at the University of California, San Francisco (supported by NIGMS P41 GM103311). The Gonen laboratory is funded by the Howard Hughes Medical Institute. Atomic coordinates and structure factors for drug-free CTD-SP1 and BVM-bound CTD-SP1 have been deposited in the Protein Data Bank with accession codes xxxx and yyyy, respectively. The maps for drug-free CTD-SP1 and BVM-bound CTD-SP1 have been deposited in the EM Data Bank with accession codes x’x’x’x’ and y’y’y’y’, respectively. The diffraction images have been deposited with the Structural Biology Data Grid under doi zzzzz. Correspondence and requests for materials should be addressed to M.Y.

## AUTHOR CONTRIBUTIONS

M.Y. and T.G. designed the experiments. J.C. purified the protein, and M.D.P. and J.C. grew the crystals. D.S. recorded the MicroED data. M.D.P. and J.H. solved the structures. M.D.P. and M.Y. analysed the structures and wrote the manuscript, assisted by T.G.

## COMPETING FINANCIAL INTERESTS

The authors declare no competing financial interests.

**Table 1.**
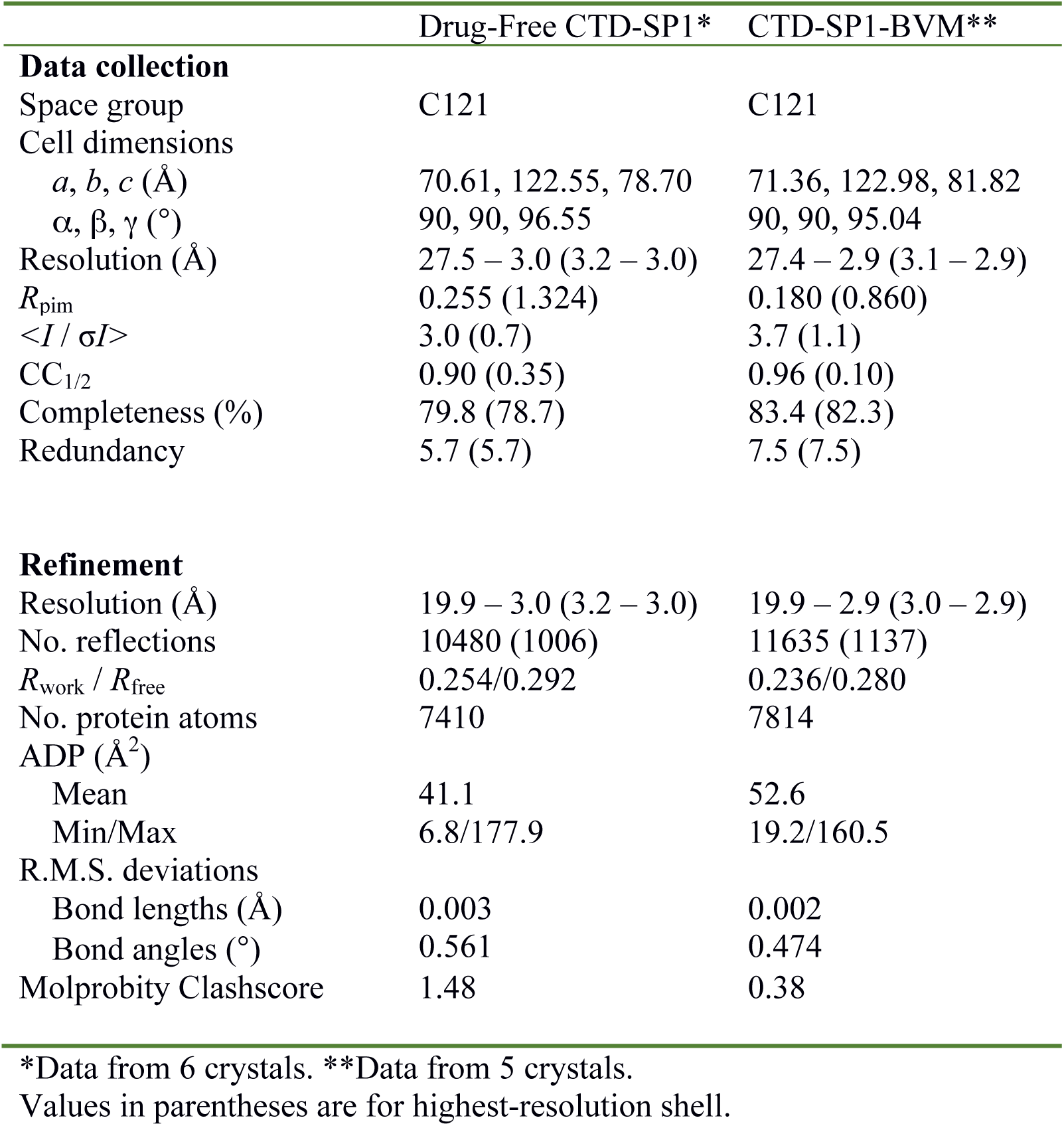
MicroED data collection and refinement statistics

## SUPPLEMENTARY METHODS

### Crystallization of CTD-SP1 and CTD-SP1-BVM

The conditions for growing microcrystals were adapted from the conditions to grow larger crystals for X-ray crystallography^17^. The crystallization solution contained 0.1 M Bis-Tris propane pH: 7.0, 1.1 − 1.2 M LiSO_4_, 0.8 − 1.0 mg/mL Gag CTD-SP1. The drug molar excess was 48x, assuming 1 BVM molecule/Gag hexamer. 600 μl of the crystallization solution was added to the well of a hanging drop plate. The protein solution was mixed with BVM (e.g. for a 1:8 protein:drug ratio, 10 μl of 1 mg/ml protein were mixed with 10 μl of a solution of 0.64 mM BVM in DMSO/water). An aliquot (1 μl) of the protein/BVM solution was mixed with 2 μl of the well solution. The crystallization plates were incubated at 17° C, and plate-like microcrystals appeared after 1-2 days. The same conditions were used to grow drug-free crystals. Initial screening of the crystals was performed by applying a 3 mm continuous carbon-coated EM grid to the crystallization drop for 90 sec, blotting the grid with filter paper and allowing the grid to dry at room temperature. Successful transfer of the crystals to the grid was first confirmed by light microscopy at 40x magnification (**Supplementary Fig. 1**). The grid was then examined in an FEI TF20 electron microscope operating at 200 kV. The unstained crystals were sufficiently thick to be readily observed from 10K-40K magnification, and successful crystallization was confirmed by electron diffraction.

### MicroED sample preparation, data collection and processing

Quantifoil EM grids were glow-discharged for 30 sec on both sides, and the grid was placed onto the crystallization droplet with the carbon-film side facing the droplet. After a 5-10 min incubation, tweezers were used to separate the grid from the droplet, and an additional 1.5 μl of solution from the same droplet was pipetted onto the back side of the grid. The grid was then mounted in an FEI Vitrobot, which was used to plunge-freeze the grid into an ethane slush, with the following settings: blotting time 7-15 sec, force 24 and humidity 40%. The grid was transferred to a Gatan 626 cryostage, and continuous rotation MicroED data were recorded using an FEI TF20 microscope equipped with a field emission electron source and a bottom-mounted TVIPS TemCam-F416 CMOS camera operating in rolling shutter mode ^20^. Diffraction patterns were recorded using 200 keV electrons with a camera length of 2 m. The electron dose rate was ~0.01 e^-^/Å^2^/sec, and the crystals were continuously rotated uni-directionally at a rate of 0.062 degrees/sec. 8 sec frames were recorded during the movie, and data typically spanned a tilt range of 0 to 45-60 degrees. Diffraction images were corrected to account for negative pixel values^27^, then they were indexed and integrated in MOSFLM ^28^. Integrated intensities from 6 and 5 crystals for the drug-free and BVM-bound samples, respectively, were scaled and merged in AIMLESS^29^.

### MicroED structure determination and map calculation

Phases for the BVM-bound and drug-free MicroED reflections were determined by molecular replacement with Phaser^30^ using the X-ray structure of hexameric CTD-SP1 (PDB 5I4T) as the search model. Cycles of real space and reciprocal space refinements were performed with Coot^31^ and phenix.refine, using electron scattering factors^32^. Torsion-angle NCS restraints were used in the early rounds of refinement. In solving the structure of CTD-SP1-BVM, the drug was never included in the model during refinements that were used for map calculations. Maximum entropy maps were calculated in Phenix and showed plausible improvements in map detail, and thus were used to aid the interpretation of the maps.

We also solved the structure of CTD-SP1-BVM independent of the CTD-SP1 X-ray structure. We generated an all-atom hexameric model of the CTD domain by combination of the X-ray crystal structure of the Gag CTD dimer (PDB 4COP^33^) and the backbone model of the CTD hexamer from the Gag CA structure determined by a combination of tomography and helical reconstruction. We aligned a copy of the 4COP monomer to each protomer of the 4D1K ^33^ CTD hexamer and performed molecular replacement with Phaser using the all-atom hexamic CTD as the search model. The top Phaser solution (TFZ = 9.4) showed well-defined density for the CTD and clear α-helical density for SP1. We generated an ideal helix in Coot within the best SP1 density and then generated the initial 6-helix bundle by application of 6-fold NCS. We then performed rounds of building and refinement in Coot and Phenix. The resultant refined BVM-bound CTD-SP1 model and associated maps recapitulated those resulting from molecular replacement with the CTD-SP1 X-ray structure.

### Electron scattering form factors and Coulombic potential maps

The presence of carboxylic acid groups at each end of BVM makes consideration of the physics of electron scattering relevant to our analysis of BVM binding to Gag, because carboxylates frequently have diminished density in maps derived from cryoEM ^34-37^ Electron diffraction is a result of elastic scattering due to Coloumb forces between the incident electron beam and the crystal, and therefore is dependent on the charge distribution in the crystal (**Fig. 2a**). Unlike atomic scattering factors for X-rays, electron scattering factors for some atoms are strongly dependent on scattering angle. In the case of negatively charged oxygen (O^-^), the atomic scattering factor is negative at low resolution (< 5.5 Å) and positive at higher resolution. This phenomenon results in a negative contribution to acidic side chains in MicroED maps calculated using low-resolution intensities (Fig. S3b,d)^37^. In contrast, maps calculated only with data in the resolution range in which scattering factors for O^-^ are positive are unaffected (**Fig. 3c,e**). The electron scattering factor for uncharged oxygen is fairly constant; therefore, neutralization of acidic side chains by direct interaction with, or proximity to, basic residues eliminates or minimizes the resolution dependence of the electron density of these side chains (**Supplementary Fig. 3b,c**). Coulomb potential neutralization of the BVM dimethylsuccinyl carboxylic acid by nearby lysine residues (K359) likely accounts for the presence of density for this moiety.

The absence of acidic side chain density can also be due to mobility and/or radiation damage. O^-^ is more susceptible to radiation damage by electrons than neutral and positively charged atoms, as shown in the analysis of radiation sensitivity of acidic side chains in single-particle cryoEM experiments^24^. This study also demonstrated that radiation damage to acidic side chains is dose-dependent, and at low total electron doses (< 10 e^-^/Å^2^) side chain density for acidic residues is not substantially diminished. The total electron dose in the CTD-SP1 MicroED experiments was 7-10 e^-^/Å^2^, and therefore, we conclude that radiation damage to acidic side chains occurred but was minimal. Future systematic studies will have to be conducted to conclusively determine the extent of radiation damage in MicroED experiments.

**Supplementary Figure 1.**
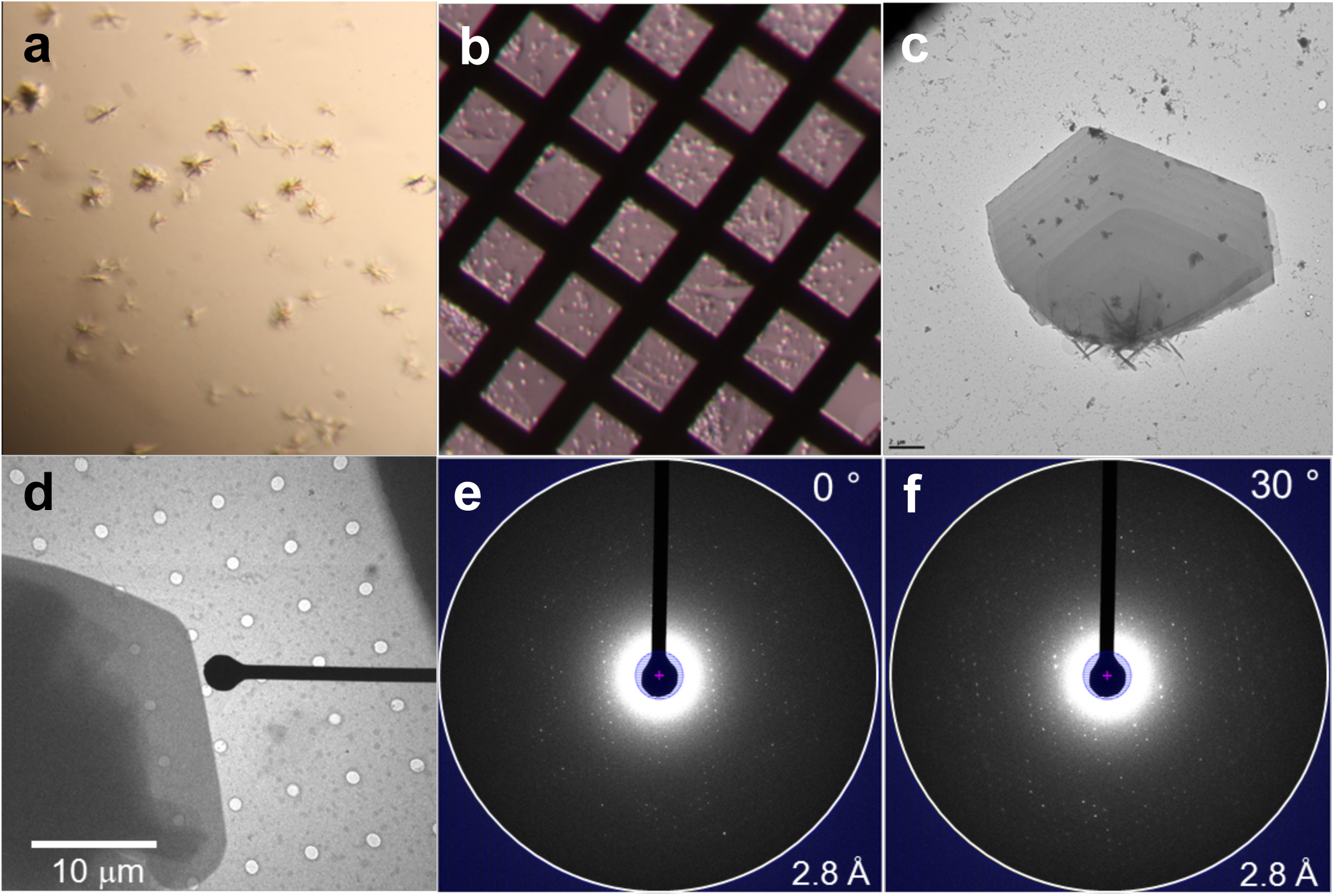
CTD-SP1 crystals and electron diffraction. **(a)** Similar to X-ray crystallography, conditions were screened to grow 3D crystals. **(b)** Crystals were transferred by placing the EM grid directly on a ~1.5 μL crystallization drop. Success of crystal transfer was assessed by light microscopy. **(c)** The 5-15 micron wide crystals were sufficiently thin that they could be imaged by EM without staining. **(d)** Diffraction movies were collected from individual, frozen-hydrated 3D microcrystals. Electron diffraction intensities **(e)** 0° and **(f)** 30° tilt) were indexed, integrated, and merged using MOSFLM. Phases were determined by molecular replacement. The programs Coot and phenix.refine were used for model building and refinement.

**Supplementary Figure 2.**
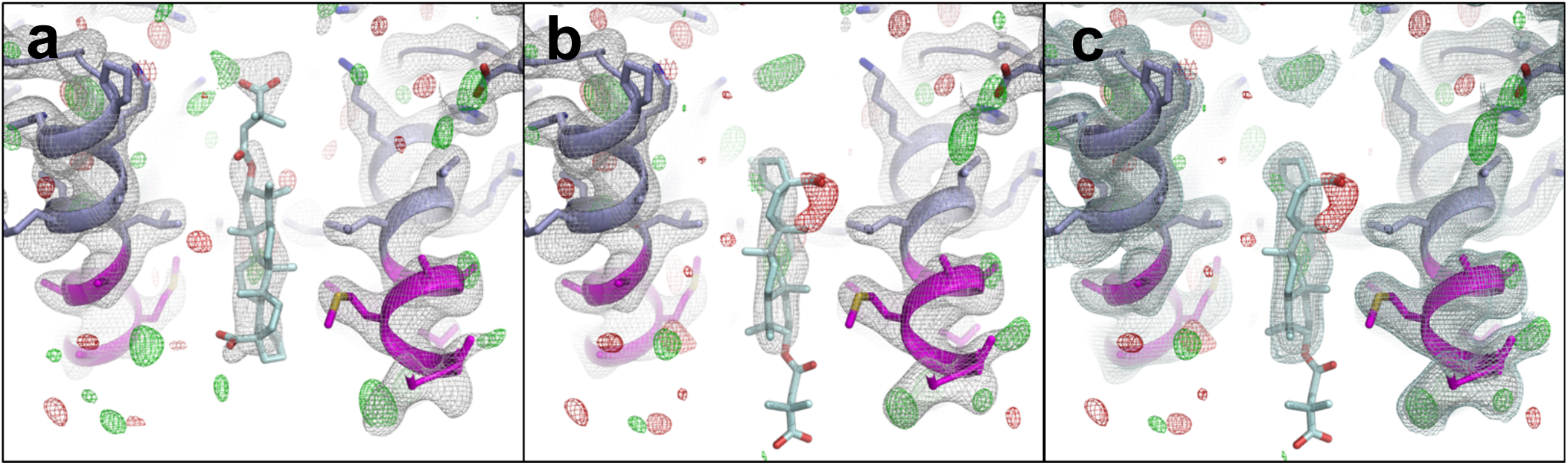
MicroED map of CTD-SP1-BVM suggests a mode of drug binding. (**a** and **b**) CTD-SP1-BVM *2Fo – Fc* (gray, contoured at 1.0 σ) and *Fo – Fc* (green/red, contoured at ±3.0 σ) maps with BVM modeled in different axial orientations. **a**, in the “up” orientation, the dimethylsuccinyl group points towards the CTD; **b**, in the “down” orientation it points away from CTD. **a**, In the “up” orientation, the pentacyclic triterpenoid moiety occupies the large map feature near the CTD-SP1 protease cleavage site, and the carboxyl of BVM is proximal to the ring of K359 lysines. **b**, In the “down” orientation, when the pentacyclic triterpenoid is positioned in the primary drug density, there is no density for the dimethylsuccinyl group and the C28 carboxyl is positioned far from K359. During refinement, without BVM, some maps included weak extensions of the primary drug density peak (> 0.6 σ) out of the helical bundle, that is, away from the CTD. However, in most maps, the drug density terminates abruptly. Following refinement with BVM in the down orientation, no density is present for the dimethylsuccinyl, which could be attributable to high mobility. **c**, Maps are the same as **b** with an additional *2Fo – Fc* contour level at 0.6 σ above the mean.

**Supplementary Figure 3.**
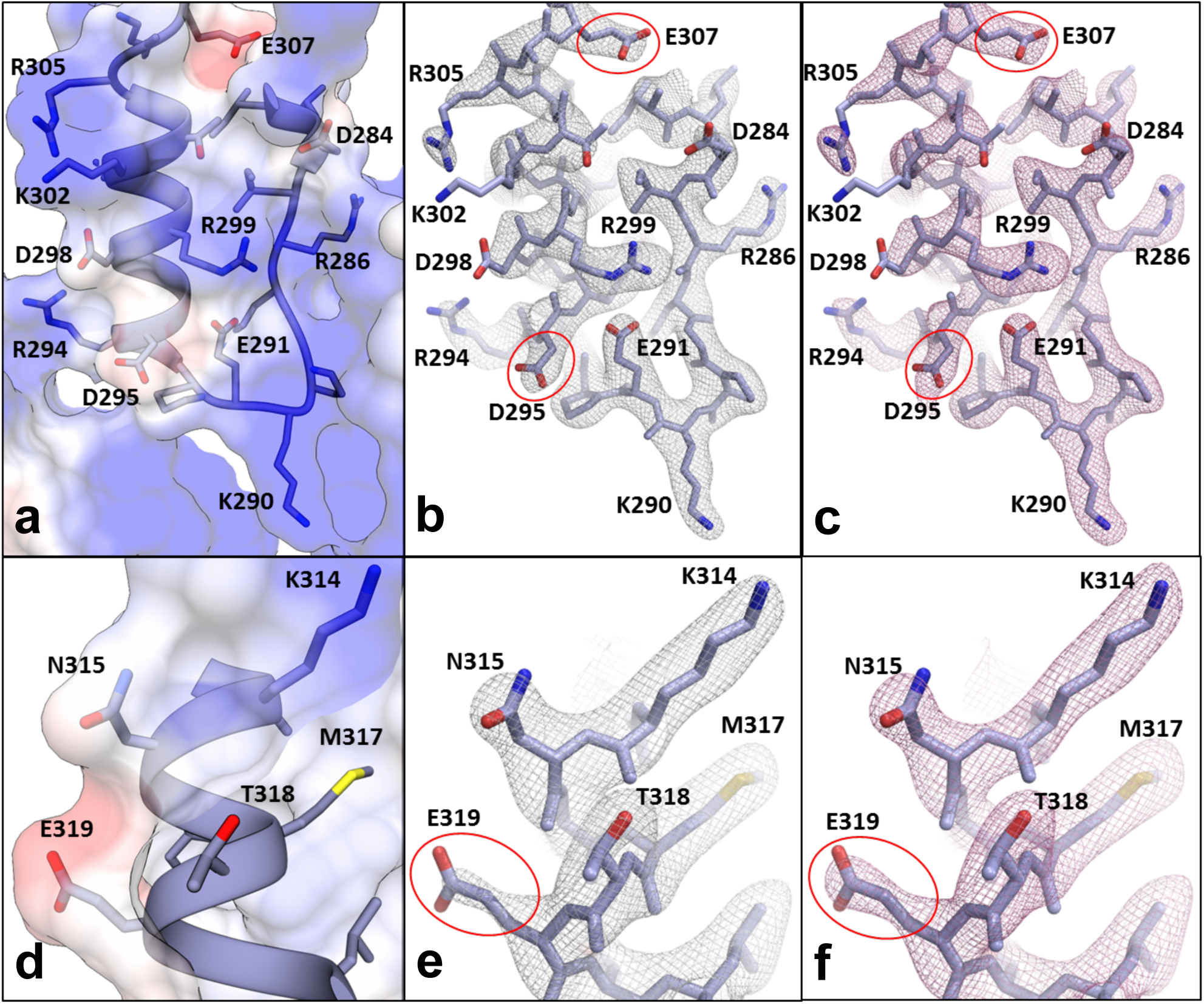
Coulomb potential map of CTD-SP1-BVM at 2.9-Å resolution. **a** and **d**, The CTD-SP1-BVM Coulomb potential surface [-8 (red) to 8 (blue) (kcal/mol*e^-^)] calculated from the model ^2^, shows significant negative charge for solvent-exposed acidic residues (D295, E307, and E319) and neutralization in the case of an acid residue involved in a salt bridge (E291). **b**, **c**, **e**, and **f**, CTD-SP1-BVM MicroED maps showing acidic and basic residues in a variety of environments. Electron atomic scattering factors are strongly dependent on charge at low resolution, and excluding these reflections results in additional side chain density for residues containing O^-^ (D295, E307, E319) **b** and **e**, 20 – 2.9 Å (gray mesh) and **c** and **f**, 8 – 2.9 Å (**pink mesh**). Scattering factors for neutral oxygen are much less dependent on resolution and are positive at all scattering angles; therefore, densities are nearly identical for acidic side chains involved in salt bridges (e.g., E291 in **b** and **c**) or neutral oxygen-containing side chains (e.g., N315 and T318 in **e** and **f**) whether the full or high resolution range is used in map calculations.

**Supplementary Figure 4.**
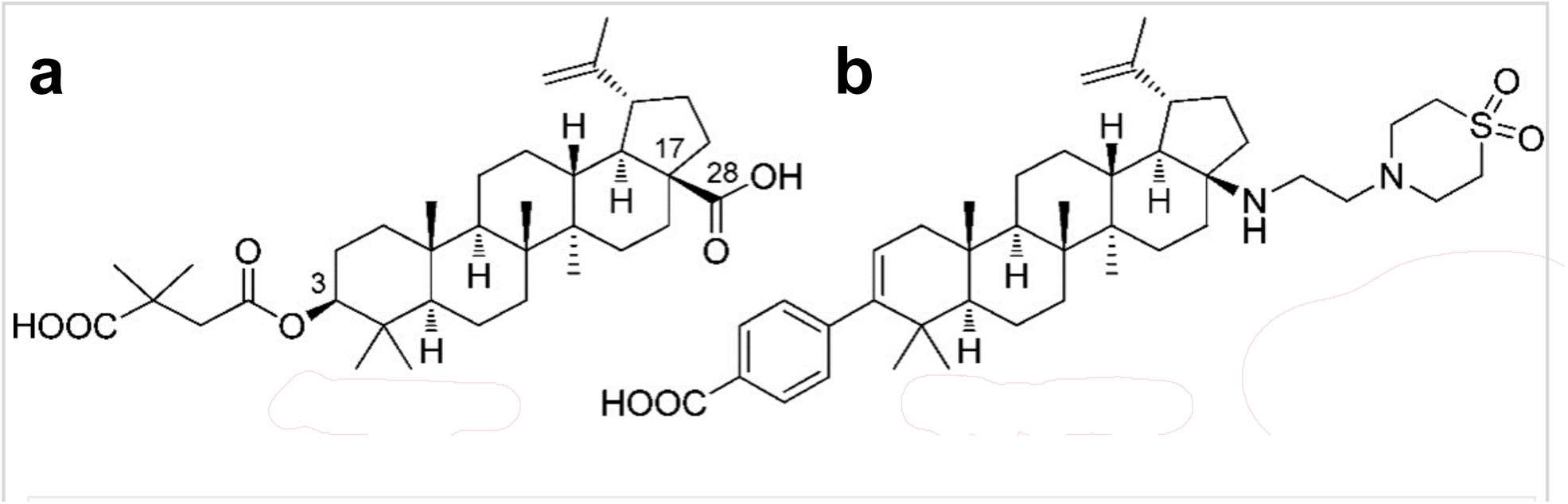
Schematic chemical representations of **a** BVM and **b** BMS-955176, a second-generation maturation inhibitor. BMS-955176 preserves the terminal carboxyl of BVM and includes a C28 extension that may provide additional stability to the 6-helix bundle in the case of resistance mutations. Adapted from^1^.

**Supplementary Figure 5.**
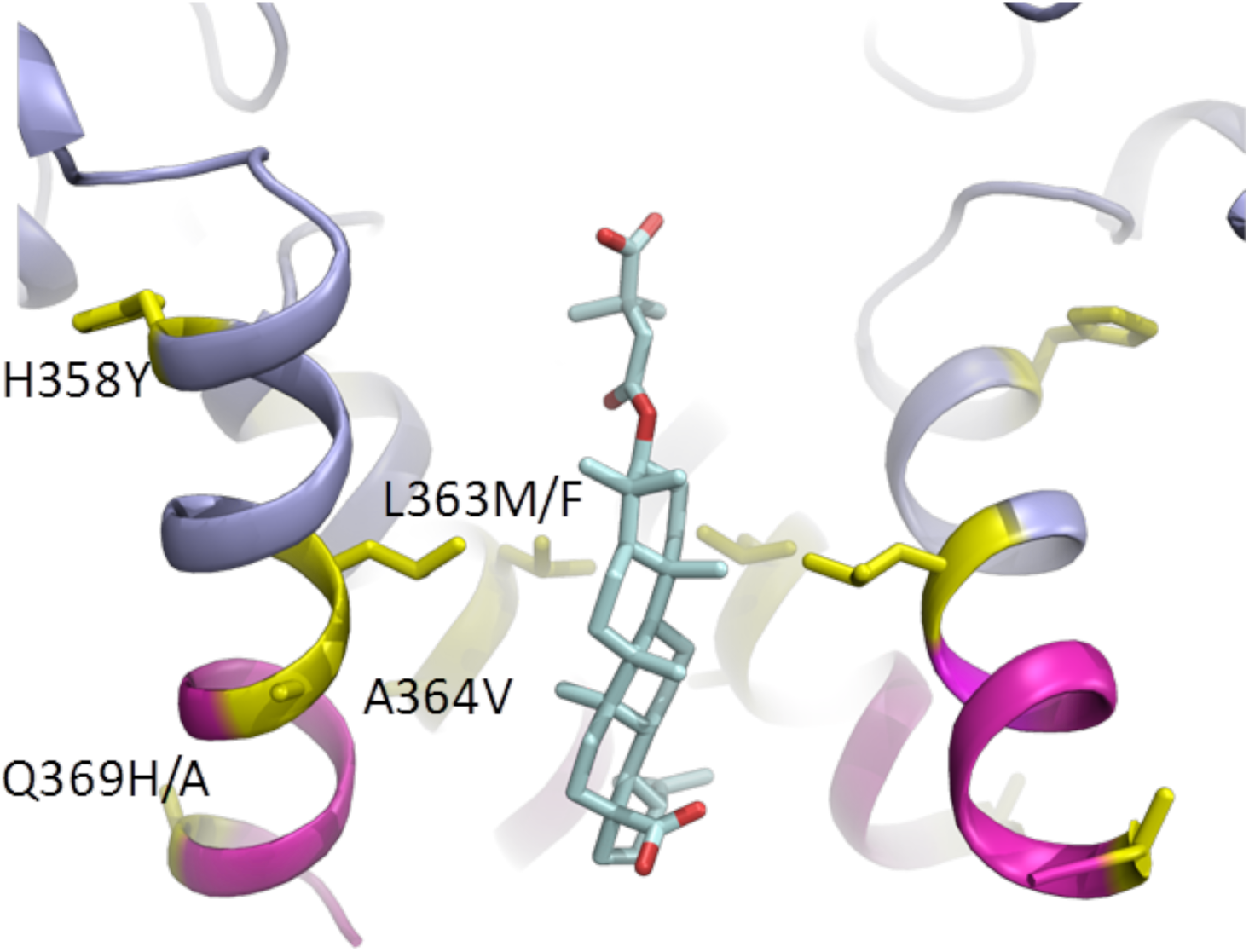
BVM resistance mutations and polymorphisms. Sites of Gag BVM resistance mutations (yellow) are shown in the context of the BVM-bound WT structure (H358Y, L363M/F, A364V, Q369H/A, V370A/M).

